# *Tbx1* and *Foxi3* genetically interact in the third pharyngeal pouch endoderm required for thymus and parathyroid development

**DOI:** 10.1101/378182

**Authors:** Erica Hasten, Bernice E Morrow

**Affiliations:** Department of Genetics, Albert Einstein College of Medicine, Bronx, NY 10461

**Keywords:** Pharyngeal apparatus segmentation, *Foxi3*, *Tbx1*, pharyngeal endoderm

## Abstract

The mechanisms required for segmentation of the pharyngeal apparatus to individual arches are not precisely delineated in mammalian species. Here, using conditional mutagenesis, we found that two transcription factor genes, *Tbx1*, the gene for 22q11.2 deletion syndrome and *Foxi3*, genetically interact in the third pharyngeal pouch endoderm for thymus and parathyroid gland development. We found that *Tbx1* is autonomously required for the endoderm to form a temporary multilayered epithelium while invaginating. E-cadherin for adherens junctions remains expressed and cells in the apical boundary express ZO-1. *Foxi3* is required autonomously to modulate proliferation and promote later restoration of the endoderm to a monolayer once the epithelia meet after invagination. Completion of this process cooccurs with expression of Alcam needed to stabilize adherens junctions and extracellular, Fibronectin. These processes are required in the third pharyngeal pouch to form the thymus and parathyroid glands, disrupted in 22q11.2 deletion syndrome patients.

## Introduction

The pharyngeal apparatus (PA) is an evolutionarily conserved structure that forms in early vertebrate embryos. The PA develops as a series of bulges, termed arches, found on the lateral surface of the head region of the embryo. During mammalian development, five pairs of pharyngeal arches numbered PA1, PA2, PA3, PA4, and PA6 (the fifth PA is transient) form subsequently, over time, from the anterior to the posterior part of the embryo (Graham, 2001a). The formation of each arch is referred to as pharyngeal segmentation. Each arch contributes to different craniofacial muscles, nerves and skeletal structures. PA1 contributes to the skull, incus and malleus of the middle ear, jaw, nerves and muscles of mastication. PA2, contributes to the skull, stapes in the middle ear, facial muscles, jaw, and upper neck skeletal structures. In addition to skeletal structures, muscles and nerves, PA3 is required to form the thymus and parathyroid glands. In mouse embryos, the thymus and parathyroid are derived only from PA3, but in humans the inferior parathyroid is derived from PA3 and PA4 contributes to the superior parathyroid gland. PA4 and PA6 contribute to the aortic arch and arterial branches (Frisdal and Trainor, 2014; Grevellec and Tucker, 2010).

Each arch is surrounded by endoderm and ectoderm derived epithelial cells forming pharyngeal pouches and clefts, respectively. Mesoderm and neural crest derived mesenchyme cells occupy the center of each arch (Graham, 2001a; Xu et al., 2005). Normal PA development is dependent on the pharyngeal endoderm (PE) cells that are needed to invaginate and promote segmentation to individual arches. The endoderm receives and sends signals from the mesenchyme to initiate morphogenesis and epithelial cells invaginate (Graham, 2008). Once each arch segments, proper patterning is also required to form derivative structures. The PE sends distinct signals in each arch to promote normal patterning (Chojnowski et al., 2014; Crump et al., 2004a; Veitch Emma, 1999; Wendling Olivia, 2000). Abnormal PA segmentation or patterning during development will lead to defects within the later structures derived from the PA. Few studies in mammals have been done on high resolution to elucidate the factors needed for segmentation.

One particular gene important for PA segmentation is *Tbx1*, encoding a T-box transcription factor implicated in 22q11.2 deletion syndrome (DiGeorge syndrome [MIM# 188400]; velo-cardio-facial syndrome [MIM# 192430]). *Tbx1*^−/−^ homozygous null mutant mouse embryos die at birth with hypoplastic and intermittent missing craniofacial muscles (Kelly et al., 2004), absent thymus and parathyroid glands, as well as a persistent truncus arteriosus (PTA) with a ventricular septal defect (VSD) (Jerome and Papaioannou, 2001; Lindsay et al., 2001; Merscher et al., 2001). *Tbx1*^−/−^ embryos have a normal PA1 but the distal arches fail to become segmented, thereby explaining, in part, why the PA derived structures are malformed (Jerome and Papaioannou, 2001; Lindsay et al., 2001; Merscher et al., 2001). *Tbx1* is expressed in the mesoderm of the head region in early mouse embryos and then throughout the endoderm, mesoderm, and distal ectoderm of the PA, while each arch forms, until mouse embryonic day (E)10.5 (Chapman et al., 1996; Garg et al., 2001; Lindsay et al., 2001). Tissue specific inactivation of *Tbx1* has been performed using the Cre-loxP system (Arnold et al., 2006a; Arnold et al., 2006b; Jackson et al., 2014; Racedo et al., 2017; Xu et al., 2005). It was found that *Tbx1* is required in all three tissues for the development of the derivative organs affected in the null mutant embryos (Arnold et al., 2006b; Choe and Crump, 2014; Racedo et al., 2017; van Bueren et al., 2010). While the formation of PA1 is normal, PA2 is hypoplastic or absent, and the distal pharyngeal arches, PA3-6 do not form (Jerome and Papaioannou, 2001; Lindsay et al., 2001; Merscher et al., 2001). Since the PE is critically important for segmentation of the distal PA (Quinlan et al., 2004; Shone and Graham, 2014), it is important to understand the genes and processes that might act downstream.

Another gene that has been shown to be important for normal PA development is *Foxi3*, which encodes a forkhead box (Fox) transcription factor. *Foxi3* is expressed in the ectoderm in the head region early in embryonic development and is then expressed in the PE and ectoderm of the PA from around the same time as *Tbx1* is expressed (Ohyama and Groves, 2004). *Foxi3* is important for epithelial differentiation within the epidermis (Shirokova et al., 2013) and has been identified in several hairless dog breeds where dogs have hair follicle and teeth defects (Drogemuller, 2008). Global null, *Foxi3*^−/−^ mouse embryos fail to form PA pouches and this results in failed PA segmentation leading to severe defects in the skull, jaw, and ears (Birol et al., 2016; Edlund et al., 2014; Khatri and Groves, 2013). It has been shown that *Foxi3* may have a cell non-autonomous effect on craniofacial neural crest cell survival because these cells undergo apoptosis in the mutant embryos (Edlund et al., 2014).

In this study, we tested whether there is a genetic interaction between *Foxi3* and *Tbx1* during mouse embryonic development. We discovered that these two factors interact specifically in the third pharyngeal pouch endoderm, needed to form the thymus and parathyroid glands. We went on to show the cellular mechanism by performing tissue specific inactivation studies. These processes require *Tbx1* function in promoting cell invagination and stratification of the PE where the arch will form, and *Foxi3* function in promoting cell invagination and restoring the endoderm to an intercalated monolayer as the final step in arch formation that is crucial for normal embryonic development.

## Results

### *Tbx1* acts upstream of *Foxi3* in pharyngeal pouch formation

Since loss of *Tbx1* or *Foxi3* disrupt the segmentation of the PA, we tested whether *Foxi3* might act upstream or downstream of *Tbx1*. Whole mount *in situ* hybridization (WMISH) using an mRNA *Foxi3* probe on *Tbx1*^−/−^ mouse embryos and wildtype (WT) littermate controls at E9.5, revealed that *Foxi3* expression was reduced in null embryos (Figure 1A and 1B). *Foxi3* is normally expressed throughout the epithelia of the PA, but in *Tbx1*^−/−^ embryos, *Foxi3* expression is visible but strongly reduced in PA1 and absent in the posterior pharyngeal region (Figure 1B). To determine if *Tbx1* expression is affected in *Foxi3*^−/−^ embryos, WMISH using a *Tbx1* probe on *Foxi3*^+/−^ control littermate and *Foxi3*^−/−^ mouse embryos showed that the *Tbx1* expression pattern was maintained in the pharyngeal mesoderm and endoderm despite the lack of segmentation of the PA (Figure 1C-1F). This indicates that either *Foxi3* acts downstream of *Tbx1* in the same genetic pathway and/or that the cells expressing *Foxi3* were lost in the *Tbx1* null mutant embryos.

**Figure 1:**
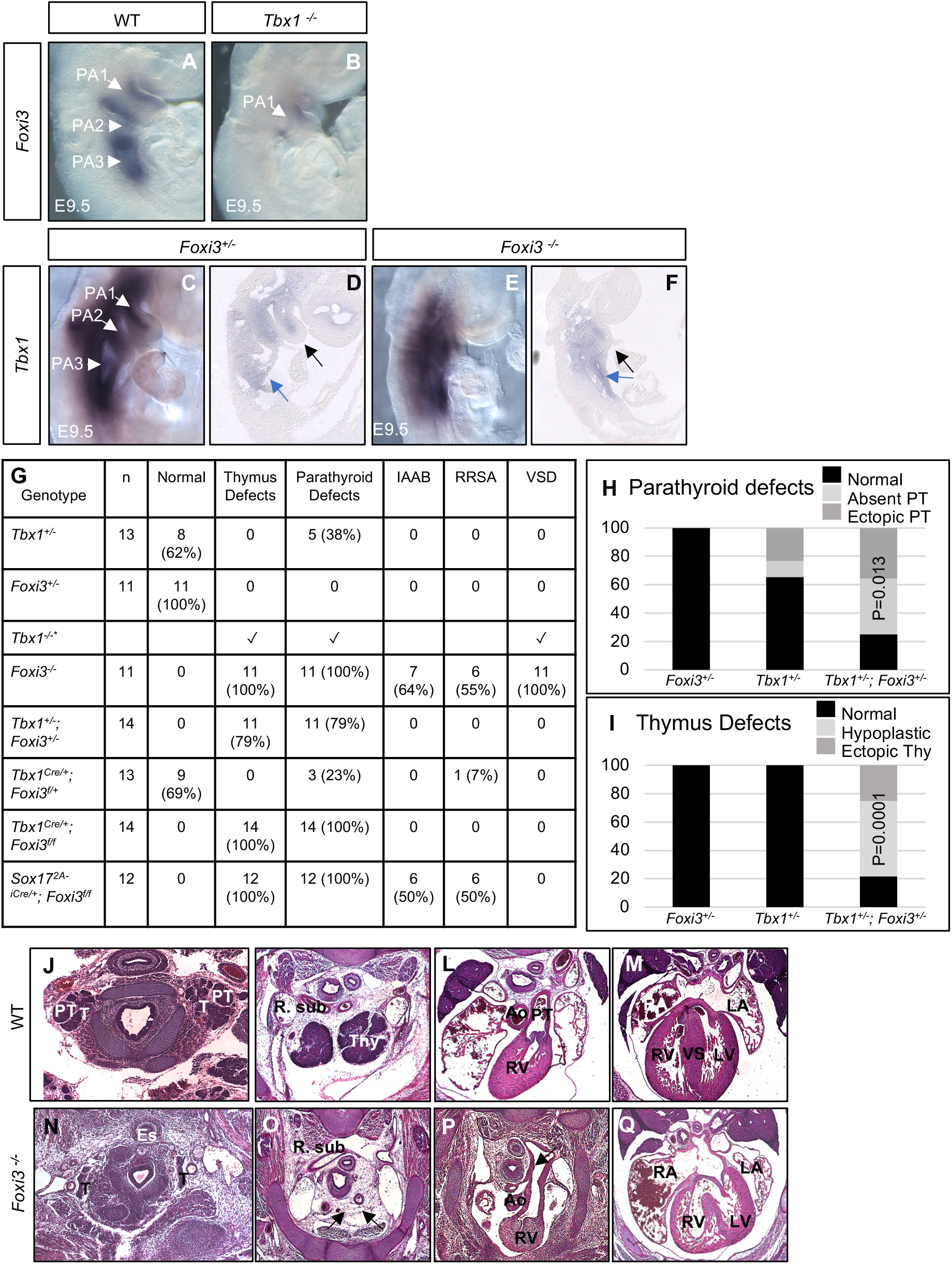
*Tbx1* and *Foxi3* genetically interact for thymus and parathyroid development. (A-F) Whole mount *in situ* hybridization (WMISH) using an antisense *Foxi3* probe on wildtype (WT) controls (A) and (B) *Tbx1*^−/−^ mouse embryos at E9.5 (n=3). (C-F) WMISH using an antisense *Tbx1* probe on whole mount and sagittal sections of control (Foxi3^+/−^) (C-D) and *Foxi3*^−/−^ (E-F) mouse embryos at E9.5 (n=3). Black arrow indicates core mesoderm, and blue arrow indicates endoderm. (G) Table summarizing defects found in each embryo with the genotype listed in the first column on the left. The second column indicates the number of embryos. The rest of the columns indicate the number or percent (in parentheses) with various defects as deterimed by histological anlaysis at E15.5. Thymus defects in *Tbx1*^−/−^ and *Foxi3*^−/−^ embryos include absent thymus and parathyroid glands. In both *Tbx1* and *Sox17* mediated *Foxi3* conditional mutants, the thymus and parathyroid glands were absent. Abbreviations: Ventricular septal defect (VSD), interrupted aortic arch type B (IAAB), retro-esophageal right subclavian artery (RRSA). Examples of defects are in Figure 1D-K. (H-I) Bar graphs summarizing parathyroid (H) and thymus (I) defects found in *Tbx1*^+/−^, *Foxi3*^+/−^, and *Tbx1*^+/−^; *Foxi3*^+/−^ embryos. Fisher’s exact two tailed test was used to determine significance between defects observed in *Tbx1*^+/−^ and *Tbx1*^+/−^;*Foxi3*^+/−^ embryos. Defects observed in *Tbx1*^+/−^;*Foxi3*^+/−^ embryos include ectopic and hypoplastic thymus and absent and ectopic parathyroid glands. (J-Q) includes histological sections of control (WT) embryos (J-M) and *Foxi3*^−/−^ embryos (N-Q) at E15.5. Arrows in (O) show absent thymus glands and in (P) shows interrupted aortic arch type B. Abbreviations: right subclavian artery (R. sub), thymus (Thy), aorta (Ao), pulmonary trunk (PT), right atrium (RA), left atrium (LA), right ventricle (RV), left ventricle (LV), ventricular septum (VS). More examples of defects that occurred in other mutant embryos are in Figures S1 and S2.

### Inactivation of *Foxi3* leads to defects in the derivative structures from the distal pharyngeal apparatus

It has been shown that *Foxi3*^−/−^ embryos have absent jaw bones, abnormal mandible development, deformed maxilla bones, absent jugal (bony arch of zygoma, cheek bone) and smaller palatines, misshapen Meckel’s cartilage, and absent ears (Birol et al., 2016; Edlund et al., 2014). At E9.5, PA segmentation of PA2 onwards does not occur in *Foxi3*^−/−^ embryos, which is consistent with previous findings (Edlund et al., 2014) (Figure S1B). Yet, there is little known about its role in formation of later embryonic structures from the distal PA derived from PA3-6. We found that at E15.5, *Foxi3*^−/−^ embryos have absent thymus and parathyroid glands (100%; n=11), interrupted aortic arch type B (IAAB, 63%; n=7), ventricular septal defect (VSD; 100%; n= 11) and retro-esophageal right subclavian artery (RRSA, 55%; n=6) as listed in Figure 1G and shown in Figure 1J-1Q. Several *Foxi3*^−/−^ embryos had both RRSA and IAAB (18%; n=2), while the remaining had either RRSA or IAAB (81%; n=9; Figure 1G, 1J-1Q)

### *Tbx1* and *Foxi3* genetically interact in PA3 for thymus and parathyroid development

Based upon the similarities in the distal PA derived defects, and that *Foxi3* was reduced in expression in *Tbx1*^−/−^ embryos, we tested whether there could be a genetic interaction between the two genes. We first tested whether single heterozygous *Foxi3*^+/−^ (Edlund et al., 2014) or *Tbx1*^+/−^ (Merscher et al., 2001) embryos had defects. At E15.5, *Foxi3*^+/−^ embryos were normal (n = 11) and *Tbx1*^+/−^ embryos had a normal thymus gland and had ectopic or absent parathyroid glands in 38% of the embryos (n = 5 Figure 1G-1I). At E15.5, double heterozygous *Tbx1*^+/−^;*Foxi3*^+/−^ embryos had a hypoplastic and ectopic thymus (n=11, 78%) and parathyroid glands (78% n=11) and this increase is statistically significant (Figure 1H-1I). Several double heterozygous embryos had both a hypoplastic thymus and ectopic parathyroid glands in comparison to WT controls (57%; n=8; Figure 1H-1I, S2A-D).

In order to determine if these double heterozygous embryos had morphology defects within PA3, histology sections were examined at E10.5. *Tbx1*^+/−^;*Foxi3*^+/−^ embryos had morphology defects in PA3 that may contribute to the phenotypes observed at E15.5 where the third pouch appears smaller in comparison to WT control embryos (Figure S2G-H). The Chorion-specific transcription factor GCMb (*Gcm2*) and Forkhead box protein N1 (*Foxn1*) genes mark the parathyroid-fated and thymus-fated domains in the pharyngeal endoderm of PA3, respectively (Bain et al., 2016; Gordon et al., 2001; Manley and Condie, 2010). We performed RNAscope *in situ* hybridization on tissue sections with probes for *Gcm2* and *Foxn1* at E11.5 when both of these genes are expressed simultaneously. In *Tbx1*^+/−^;*Foxi3*^+/−^ embryos, expression of both markers was reduced in comparison to wildtype littermate controls (Figure S2J-K). We then examined normal PA segmentation to better understand the cellular basis of the defects that were observed.

### Process of pharyngeal segmentation in wildtype embryos

The PE and ectoderm cells, together, can be referred to as epithelial cells. The epithelial cells have an important role in PA segmentation and patterning (Campbell et al., 2011; Graham, 2001a, b);(Choe and Crump, 2015; Tsuchiya et al., 2018). Cell adhesion molecules enable cells to assemble together by connecting the internal cell cytoskeletons to determine overall architecture of tissues and intercellular signaling. E-cadherin is a cell-cell adhesion protein forming adherens junctions that bind cells tightly to each other (Reviewed in (Gumbiner, 1996). Another protein expressed within the peripheral membrane on the apical side of epithelial cells is zonula occluden-1 (ZO-1). ZO-1 proteins form permeable barriers in adherens junctions (Anderson et al., 1989).

We noticed in wildtype embryos from E8.5-E9.0, the intercalated monolayer of ectoderm and endoderm cells disappeared and a transitional multilayered or stratified epithelium appeared where invagination began (Figure 2A-2E). The two processes appear to occur simultaneously. We refer to the first step of invagination with transient stratification as the first transition of pouch morphogenesis. E-cadherin was expressed in the monolayer and multilayered epithelia, indicating that adherens junctions remained during this process (Figure 2A-2E). ZO-1 expression in the apical side of the outermost layer remained, indicating that polarity was retained (Figure 2D-2E). However, cells in the additional layers of epithelium internal to the outermost layer, did not express ZO-1, and therefore did not have a particular apical/basal polarity (Figure 2D-2E). After invagination was complete, the cells reverted back to an intercalated single layer of cells where the endoderm pouch meets the ectodermal cleft, with apical expression of ZO-1 (Figure 2F-2G). We refer to this second step of restoration of a monolayer of endoderm and ectoderm, as the second transition of pouch morphogenesis. At E10.5, PA formation was completed and each arch was surrounded by an intercalated epithelium derived from endoderm and ectoderm (Figure 2H-2J). This process continued over time for the formation of PA3, PA4 and PA6 (Figure 2H-2J).

**Figure 2:**
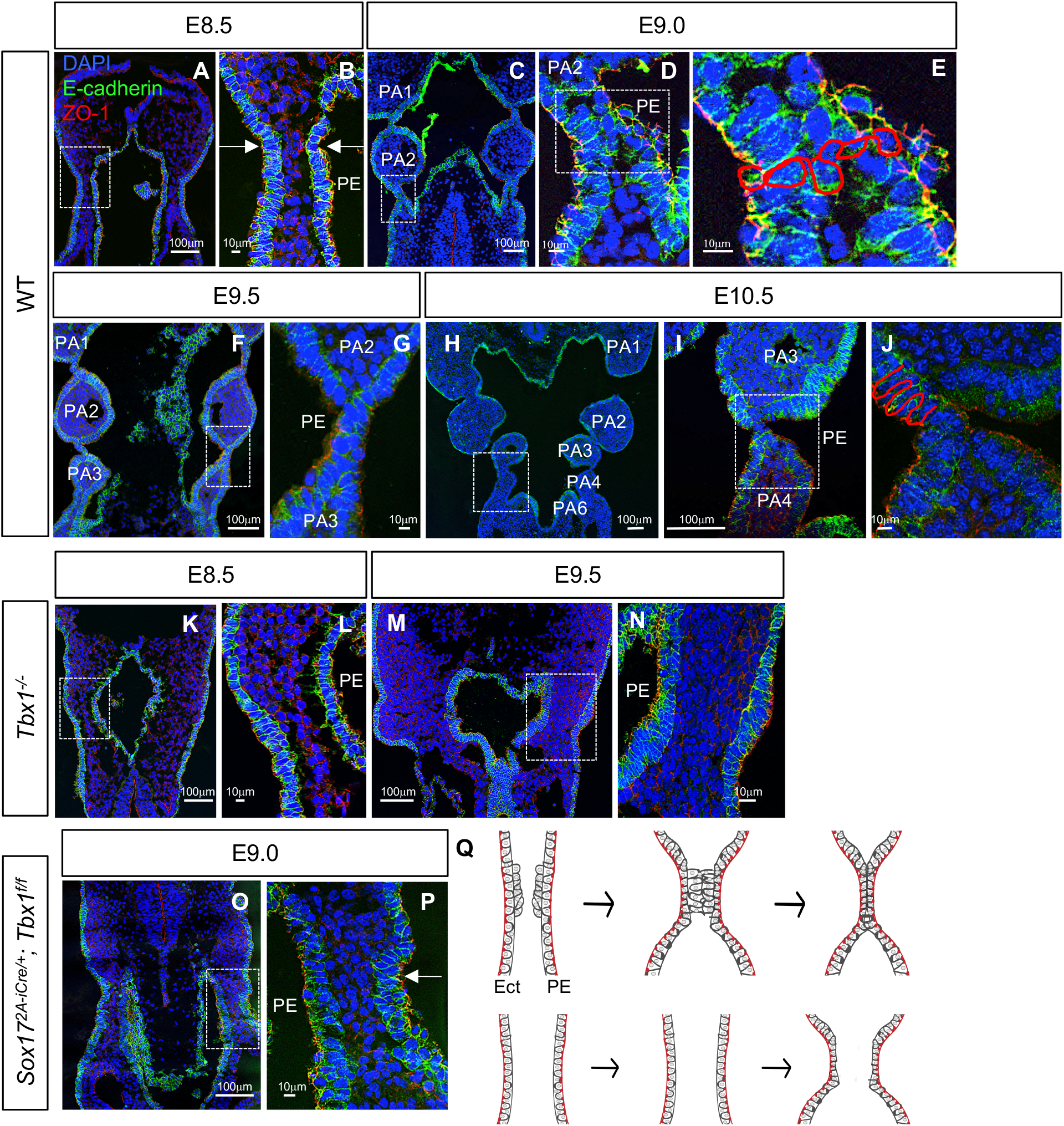
Endoderm and ectoderm undergo morphogenesis, stratification, and restoration during PA segmentation and these processes are disrupted in *Tbx1* mutant embryos. (A-I\J) To visualize epithelial cells within the PA, DAPI (blue), E-cadherin (green), and ZO-1 (red) antibodies were used on coronal WT sections at various embryonic stages from E8.5-E10.5 (n=5 for each stage). E8.5 embryos (A-B), E9.0 (C-E), E9.5 (F-G), and E10.5 (H-J) are shown. White boxes indicate where each section is magnified. PE indicates the pharyngeal endoderm for each section. Red lines trace nuclei of E-cadherin positive epithelium. (K-N) Epithelial cells in *Tbx1*^−/−^ coronal sections were visualized at E8.5 (K-L) and E9.5 (M-N) for n=3. PE indicates pharyngeal endoderm. (O-P) *Sox17^2A-iCre/+^;Tbx1^f/f^* conditional mutant embryos at E9.0 when PA2-PA3 will become segmented (n=3). (Q) Cartoon depicting the WT (top) and *Tbx1* null (bottom) PA segmentation processes. WT embryos undergo PA morphogenesis where epithelium invaginate to form each arch. Then, to create each segment, the epithelium undergoes a transient multilayering process or stratification. Finally, the epithelium is restored to a monolayer of cells where each arch segments. Segmentation fails in *Tbx1*^−/−^ embryos. The epithelia fail to invaginate and fail to form multilayers as shown in the cartoon.

### *Tbx1* in the PA endoderm initiates invagination and stratification of PE cells

Previous studies have shown that global or endoderm inactivation of *Tbx1* resulted in failed segmentation of PA2-PA6 (Arnold et al., 2006b; Jackson et al., 2014; Jerome and Papaioannou, 2001; Lindsay et al., 2001; Merscher et al., 2001). To test the mechanism by which segmentation was disrupted, *Tbx1*^−/−^ mutant embryos were generated and immunofluorescence with antibodies to E-cadherin and ZO-1 was performed on coronal tissue sections. In global null mutant embryos, invagination and stratification to form transitional multilayers was disrupted (Figure 2K-2N). This is illustrated in the cartoon on Figure 2Q, where we show epithelial cell invagination and stratification occurs in the wild type embryos during pouch and cleft formation while this fails in *Tbx1* mutant embryos. Nonetheless, E-cadherin and ZO-1 expression was normal, so that the existing cells did not lose their epithelial characteristics or apical/basal polarity (Figure 2K-2N). We then performed endoderm specific inactivation to determine whether this process was cell type autonomous. In *Sox17^2A-iCre/+^;Tbx1^f/f^* mutant embryos at E9.0, endodermal invagination and stratification was disrupted (Figure 2O and 2P). Similar to the global null mutant embryos, E-cadherin and ZO-1 expression was normal in the single layer of PE. Invagination and cell shape were normal in the ectoderm (Figure 2P). This demonstrates that *Tbx1* is required in the first transition of pouch morphogenesis, to initiate invagination and stratification within the endoderm in an autonomous manner.

### Inactivation of *Foxi3* in endodermal cell lineages causes defects in PA derived structures

The PA develops from E8.5-E10.5, in which PA3 forms by E9.5. *Tbx1* is expressed in the PE at E8.5 and E9.5 (Payen et al., 2015; Zhang et al., 2005; Zhang et al., 2006). We used RNAscope *in situ* hybridization on coronal sections from embryos to determine if there is overlap between *Foxi3* and *Tbx1* mRNA expression in wild type embryos at E9.5 (Figure 3A-3L). In PA1 and PA2, *Tbx1* and *Foxi3* expression overlaps in the endoderm but not the ectoderm or mesoderm (Figure 3A-3H). In PA3, there was co-expression throughout the endoderm and ectoderm, and the expression of both genes is strongest where the two epithelia are invaginating in the process of segmentation (Figure 3I-3L). The expression pattern of both genes at E9.5 is summarized in Figure 3M. We also evaluated the expression patterns of both genes at E10.5 when PA4 and PA6 have formed. At this stage, *Foxi3* expression was reduced throughout the PA, and *Tbx1* continued to be expressed within the endoderm and ectoderm PA3-PA4 (data not shown). The region of strongest co-expression in PA3 and absence in PA4-6, may explain the basis of parathyroid and thymus defects in double heterozygous mutant embryos and reduced expression of *Gcm2* and *Foxn1*, but no defects in the aortic arch or heart.

**Figure 3:**
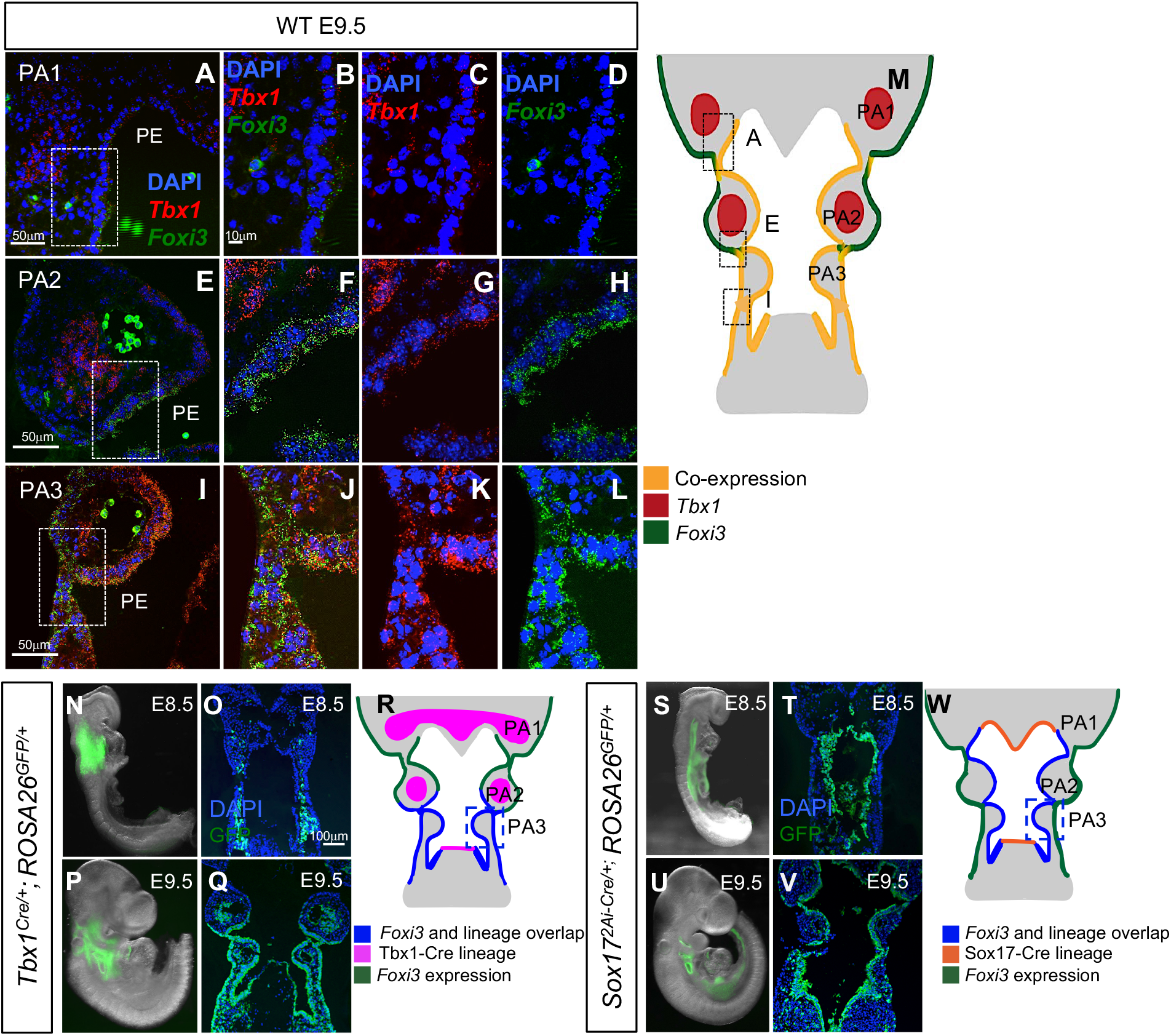
*Foxi3* and *Tbx1* expression and *Tbx1^Cre^,Sox17^2A-iCre^* lineages overlap within PA3. (A-L) RNAscope *in situ* hybridization with mRNA probes for *Foxi3* and *Tbx1* on coronal sections from a WT embryo at E9.5. *Tbx1* expression is shown in red and *Foxi3* expression is shown in green. (A-D) Image of PA1 where *Tbx1* and *Foxi3* expression overlaps in the PE. These images correspond to box A in cartoon M. (E-H) Image of PA2 where *Tbx1* and *Foxi3* are co-expressed within the PE. These images correspond to box E in cartoon M. (I-L) In PA3 there is co-expression throughout the endoderm and ectoderm especially where PA4 will segment. These images correspond to box I in cartoon M (n=4). PE indicates the pharyngeal endoderm for each section. (M) The expression pattern of both genes depicted in a cartoon of the PA at E9.5. (N-Q) Lineage tracing of *Tbx1^Cre/+^;Rosa26^GFPf/+^* embryos. GFP expression is visible in whole mount embryos and the corresponding coronal sections at E8.5 (N-O) and at E9.5 (P-Q); n=3 for each stage. (R) Cartoon depiction of where *Tbx1^Cre^* will inactivate *Foxi3* expression (blue) and where expression will not be affected (green) at E9.5. (S-W) Lineage tracing of *Sox17^2A-iCre/+^;Rosa26^GFPf/+^* embryos. GFP expression is visible in whole mount embryos and the corresponding coronal sections at E8.5 (S-T) and E9.5 (U-V); n=4 for each stage. (W) Cartoon depiction of where *Sox17^2A-iCre/+^* will inactivate *Foxi3* expression (blue) and where gene expression will not be affected (green) at E9.5.

In order to establish the role of *Foxi3* within the *Tbx1* lineage to explain the PA3 derived defects in double heterozygous embryos, *Tbx1^Cre/+^* mice were used (Vitelli et al., 2006) to inactivate both alleles of *Foxi3*. Prior to this, we first crossed *Tbx1^Cre/+^* mice with Rosa26^*GFPf/+*^ mice. GFP fluorescence was only observed in the pharyngeal mesoderm, at E8.5 (Figure 3N and 3O) and in both the pharyngeal mesoderm plus epithelia of the PA at E9.5 (Figure 3P-3R). The *Tbx1^Cre/+^* is a knock-in allele for *Tbx1* and it is therefore heterozygous for *Tbx1* (Vitelli et al., 2006). However, there is a delay in Cre recombinase activity (Figure 3N-3R) as compared to *Tbx1* mRNA expression. This timing difference may explain why *Tbx1^Cre/+^;Foxi3^f/+^* embryos at E15.5 do not exhibit a more severe phenotype than what occurs in *Tbx1*^+/−^ embryos, as compared to that in *Tbx1*^+/−^;*Foxi3*^+/−^ embryos (Figure 1G). Loss of *Foxi3* expression in PA3 of *Tbx1^Cre/+^;Foxi3^f/f^* embryos is confirmed in Figure S3.

At E15.5, *Tbx1^Cre/+^;Foxi3^f/f^* mutant embryos were compared to *Tbx1^Cre/+^;Foxi3^f/+^* controls (n=13) to determine if there were any PA derived defects. The *Tbx1^Cre/+^;Foxi3^f/f^* embryos had absent thymus and parathyroid glands (100% n=14), similar to the *Foxi3*^−/−^ embryos (Figure 1G). In order to determine if *Foxn1* (thymus) and *Gcm2* (parathyroid) expression (Gordon et al., 2001) was affected, we performed RNAscope *in situ* hybridization with probes for these genes and found the expression of both markers were greatly reduced (Figure S2). We also note that there are severe morphology defects in this region of the embryo (Figure S2). Despite having an absent thymus and parathyroid glands, these mutant embryos did not have cardiovascular defects (Figures 1G and S4). While expression of *Foxi3* was significantly reduced as determined by WMISH (Figure S3E-S3H) in conditional mutant embryos, we also tested *Tbx1^Cre/+^;Foxi3^f/−^* embryos in order to further inactivate *Foxi3* and found that the embryos lacked the thymus and parathyroid glands but similar to above, they had no intracardiac or aortic arch defects (n = 8; data not shown). Therefore, there is an interaction between *Tbx1* and *Foxi3* in the third pharyngeal pouch endoderm but not in the fourth pouch endoderm, which contributes to the aortic arch and subclavian arteries. It is also possible that the lack of aortic arch anomalies could instead be due to timing of *Tbx1* gene expression and delayed timing of Cre activity within the PE (Figure 3N-3R).

*Foxi3* is expressed in the PE and ectoderm in the PA (Figure 3 and (Ohyama and Groves, 2004). In order to determine the role of *Foxi3* within the endoderm, we performed tissue specific inactivation of *Foxi3* using the *Sox17^2A-iCre/+^* mouse (Figure 3S-3W). We first confirmed that the PE population is marked by GFP expression using *Sox17^2A-iCre/+^;ROSA26^GFP/+^* embryos (Engert et al., 2009) (Figure 3S-3W). We found that *Sox17^2A-iCre/+^;Foxi3^f/f^* embryos had a hypoplastic distal PA as compared to *Sox17^2A-iCre/+^;Foxi3^f/+^* controls (Figure 3S-3W, S1C). Recombination occurred prior to E8.5 as indicated by robust green fluorescence at E8.5 in Figure 3S-3T. We found that *Foxi3* expression was reduced in these conditional mutants using WMISH (Figure S3). The PA was not as severely affected as in the *Foxi3*^−/−^ null mutant embryos, as PA2 was present, albeit hypoplastic (Figure S1C). Nonetheless, the distal PA did not become segmented. At E15.5, these endodermal conditional loss of function mutants had absent thymus and parathyroid glands in all embryos (100%; n=12) as well as IAAB (50% n=6), and RRSA (50% n=6) in half the embryos, as shown in Figure 1G and Figure S1. As compared to *Foxi3*^−/−^ mutant embryos that always have an aortic arch defect plus VSD, in these conditional mutant embryos with IAAB, there was normal septation of the ventricles (Figure 1G and S1L). Several mutant embryos had both an RRSA and IAAB (33% n=3), while 33% (n=3) had just an RRSA and 33% (n=3) had an IAAB, whereas a few had normal aortic arches (33% n=3) (Figure 1G). India ink injections showed that the 4^th^ arch artery was absent in *Sox17^2A-iCre/+^;Foxi3^f/f^* embryos at E10.5, similar to what we observed in *Foxi3*^−/−^ embryos (Figure S5). Based upon this data, *Foxi3* expression in the ectoderm may have an additional role in the septation of the ventricles.

### Loss of *Foxi3* in the Tbx1 expressing lineage disrupts PA3 formation

In the *Tbx1^Cre/+^;Foxi3^f/f^* mutant embryos, only PA3 derivative structures were affected, being the thymus and parathyroid glands. At E8.5 the process of invagination, stratification and resolution of the epithelial populations to form the pouch and cleft of PA2 was normal (Figure 4A-4H). But, at E9.5, PA3 formation was specifically disrupted. Here, the second transition of pharyngeal pouch formation was disrupted. We found that the stratification process was exaggerated in the PE and there was failed invagination of the PE and ectoderm (Figure 4I-4P). Further, the PE and ectoderm of PA3 fails to resolve to monolayers (Figure 4M-4P) as compared to controls (Figure 4M-4P). As before, the PE of PA3 maintains its epithelial identity as marked by expression of E-cadherin (Figure 4N). The outer cell layer maintained normal apical/basal polarity since ZO-1 was expressed normally (Figure 4O). However, endodermal cells within the stratified epithelium did not express ZO-1 and therefore did not have apical/basal polarity. At E10.5, the third pouch is absent in the conditional mutants (Figure 4S and 4T) in comparison to controls (Figure 4Q and 4R). The cartoon in Figure 4Q depicts the epithelial morphology from E8.5-E9.5 in mutants and controls. Thus, the genetic interaction between the two genes is only within PA3.

**Figure 4:**
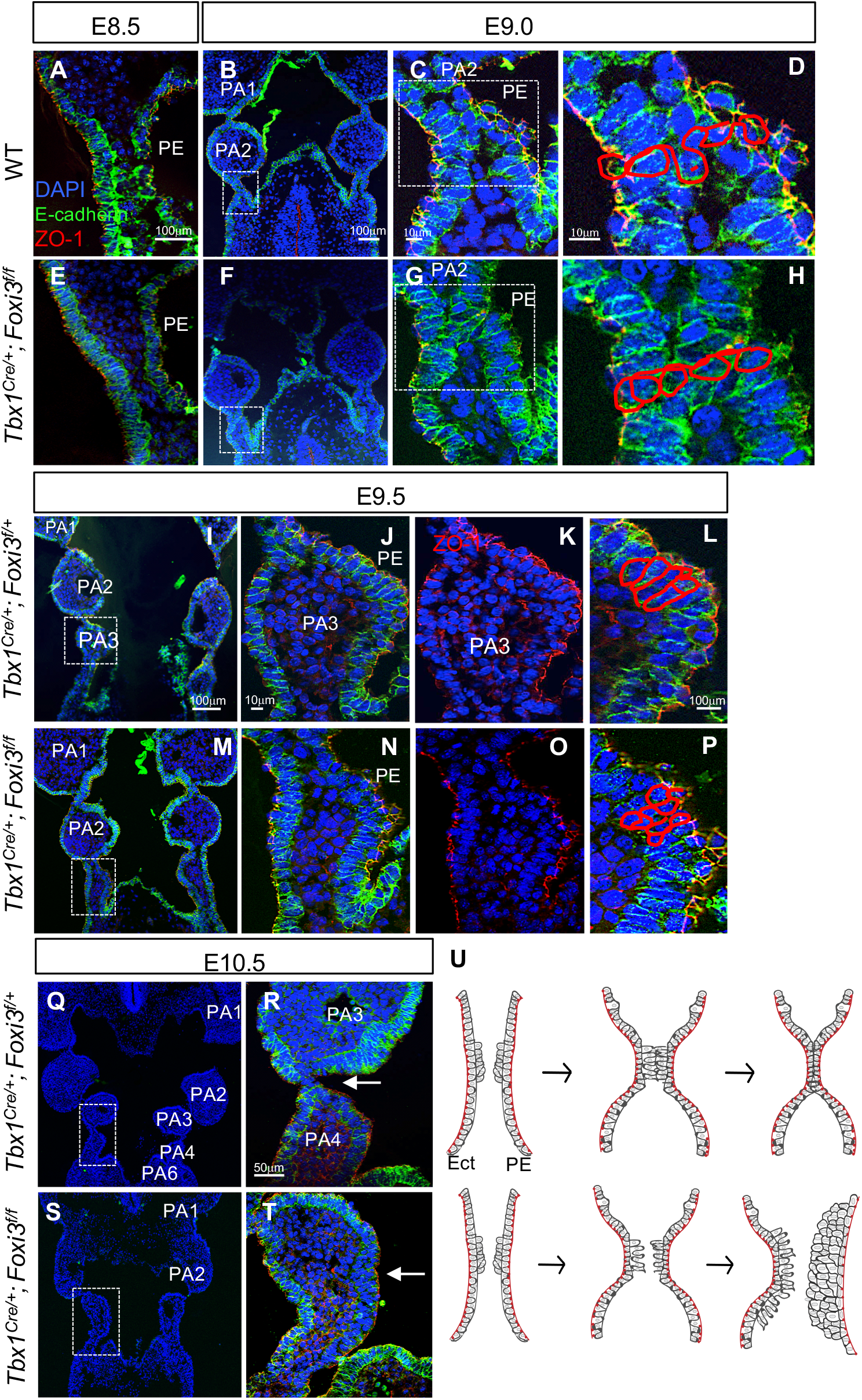
Inactivation of *Foxi3* within the *Tbx1^Cre/+^* lineage disrupts PA3 morphogenesis. (A-T) To visualize epithelial cells within the PA, DAPI (blue), E-cadherin (green), and ZO-1 (red) antibodies were used on coronal sections at E8.5-E9.0 (n=3), E9.5 (n=5) and E10.5 (n=4) in WT (A-D) or *Tbx1^Cre/+^;Foxi3^f/+^* (I-L, Q-R) controls and in *Tbx1^Cre/+^;Foxi3^f/f^* (E-H, M-P, S-T) mutants. White boxes indicate the location of the higher magnification images. Red lines trace nuclei of E-cadherin positive epithelial cells. White arrows in E10.5 images (R, T) indicate absent 3^rd^ pharyngeal pouch in mutant embryos. (Q) The cartoon depicts the epithelial morphology at these stages in controls (top) and *Tbx1^Cre/+^;Foxi3^f/f^* mutants (bottom) at E8.5-E9.5. In *Tbx1^Cre/^+;Foxi3^f/f^* embryos, morphogenesis of PA1-PA2 is normal, however it is disrupted when PA3-4 forms. When PA3-PA4 forms, the first transition is initiated in the PE and ectoderm and epithelial cells form stratified layers. However the second transition does not occur. There is excessive stratification of the epithelia and the shapes of the cells are abnormal.

### Resolution of the pharyngeal epithelia to monolayers is disrupted in *Foxi3* mutants

To test for a function of *Foxi3* in pharyngeal segmentation, we examined *Foxi3*^−/−^ embryos. We found a failure of the first transition of pouch morphogenesis. At E8.5, the epithelial cells failed to initiate invagination (Figure 5A-5D). There was also failure in the second transition of pouch morphogenesis. The endodermal cells formed excessive stratified multilayers, did not invaginate or resolve to form an intercalated monolayer in *Foxi3*^−/−^ embryos at E8.5 or E9.5 (Figure 5E-5L). The ectodermal cells formed a monolayer at these stages, however the cells were misshapen (Figures 5L). All epithelial cells expressed E-cadherin and those on the outer edge, expressed ZO-1, indicating that they did not lose epithelial identity nor polarity (Figure 5J-5L). However, cells within the stratified epithelium did not express ZO-1 (Figure 5K). As mentioned before, PA1 forms, while PA2 is hypoplastic and the distal pharyngeal arches fail to form in *Sox17^2A-iCre/+^;Foxi3^f/f^* mutant embryos. In *Sox17^2A-iCre/+^;Foxi3^f/f^* embryos, the PE cells failed to invaginate at E8.5-E9.5 (Figure 5M-5N). In contrast, the ectodermal cells were able to invaginate normally (Figure 5N). The endoderm had the same defects as in *Foxi3^−/−^ embryos, where cells formed more layers than normal and didn’t resolve to single layers (Figure 5O-5R). As in the *Foxi3*^−/−^* embryos, E-cadherin was expressed throughout the epithelia and ZO-1 was expressed primarily the outer layer of cells (Figure 5O-5R).

**Figure 5:**
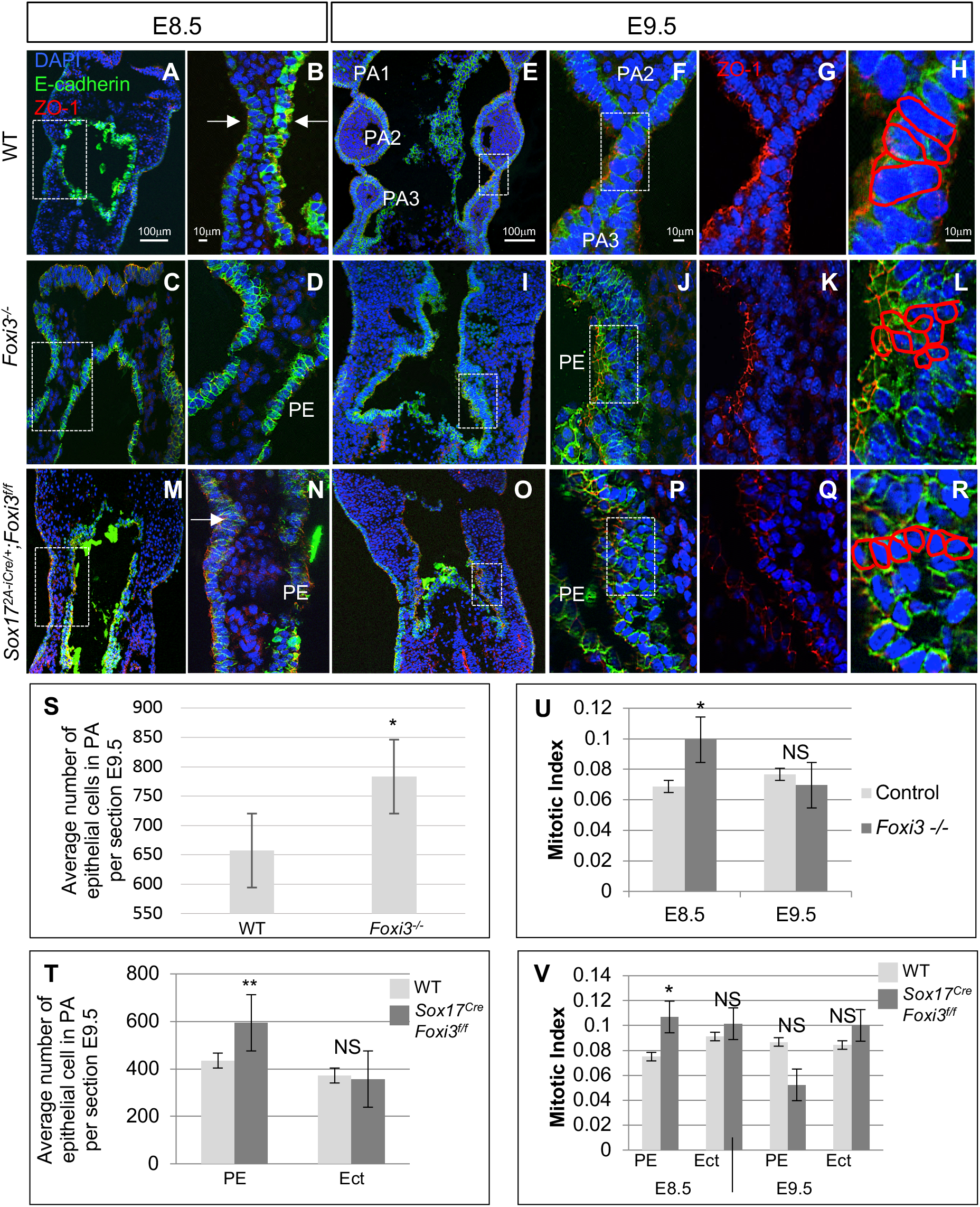
*Foxi3*^−/−^ and *Sox17-2A-iCre/+;Foxi3^f/f^* mutant embryos have defects in PA segmentation. (A-O) DAPI (blue), E-cadherin (green), and ZO-1 (red) mark epithelial cells on WT control (A-F), *Foxi3*^−/−^ (G-L) and *Sox17^2A-iCre/+^;Foxi3^f/f^* (M-R) coronal sections of mouse embryos from E8.5-E9.5 (n=4). White boxes indicate the location of higher magnification images. White arrows indicate where invagination occurs. Red lines trace nuclei of E-cadherin positive cells. (S-T) Quantification of epithelial cells within the pharyngeal apparatus at E9.5 in controls and mutants. Error bars represent standard error. One asterisk indicates P value <0.05; two asterisks indicate P value <0.005 (n=4 each). (U-V) Quantification of proliferation assays include counting E-cadherin positive epithelial cells within the PA and calculation of the mitotic index, which is the ratio of pH3 positive cells within the E-cadherin positive epithelial cells. Error bars represent standard error, and asterisks indicate P value <0.05. Pharyngeal endoderm (PE) and Ectoderm (Ect) were quantified separately in conditional mutants. At E8.5 a total of n=6 were examined for each control and mutant embryo; at E9.5, n=4 for were examined for each control and mutant embryo.

E-cadherin positive epithelial cells within the PA were quantified and there were significantly more cells within *Foxi3* mutants in comparison to controls (Figure 5S and 5T). These defects within the epithelial cells may explain why normal PA segmentation does not occur. At E10.5, the epithelial cells, despite existence of additional multilayers, and failure to return to monolayers, begin to invaginate after a long delay (Figure S6). This indicates that some of the PE is able to partially respond to molecular signals required for invagination.

### Cell proliferation increases in *Foxi3* mutant embryos at E8.5 but not at E9.5

As mentioned above, we detected an increase in the number of epithelial cells within the PA in *Foxi3*^−/−^ embryos at E9.5 (Figure 5P). There was a 40% increase in the number of endodermal cells and, but no increase was detected of ectoderm cells in the PA of *Sox17^2A-iCre/+^;Foxi3^f/f^* mutant embryos versus controls, at E9.5 (Figure 5Q). We then decided to test whether cell proliferation was increased at E8.5-E9.5. For this, pH3 antibodies was used to mark proliferating cells on serial sections on mutant versus control embryos (Figure S7). When calculating the ratio between proliferating cells versus total E-cadherin positive cells, there was a significant increase at E8.5 in *Foxi3*^−/−^ and *Sox17^2A-iCre/+^;Foxi3^f/f^* embryos (Figure 5U and 5V). At E9.5 there was no difference in proliferation between mutant and control embryos (Figure 5U and 5V). This indicates that increases of cell proliferation at E8.5 can partially explain why there is an increase of layers of epithelial cells within the PA at E9.5.

### Alcam and Fibronectin expression is less in *Foxi3* null embryos

It has been shown that activated leukocyte cell adhesion molecule (Alcam; also called CD166, Neurolin, or DM-GRASP) is an extracellular molecule that functions for neuronal cell migration, cardiac morphogenesis, and neural crest differentiation during development (Gessert et al., 2008; Lee et al., 1996; Tomita et al., 2000). Alcam has been shown to stabilize adherens junctions in epithelial cells (Tomita et al., 2000) or through physical forces to bring adjacent PE cells closer together in zebrafish pouch formation (Choe et al., 2013). To demonstrate that *Foxi3* plays a role in invagination and resolving the epithelial layers to form monolayers at E9.0, as well as to determine what downstream effectors may be important for PA segmentation that was reported in zebrafish (Choe and Crump, 2015), we performed immunofluorescence for Alcam on tissue sections from *Foxi3*^+/−^ and *Foxi3*^−/−^ embryos at E9.0 when epithelial cells in PA1-2 enter the second transition and begin to resolve to form monolayers and when epithelial cells in PA3 enters the first transition and begins to invaginate and stratify (Figure 6A-6D). In *Foxi3*^−/−^ embryos, Alcam expression remained on the basal side of the ectoderm but was strongly reduced on the basal side of the endoderm (Figure 6D). In zebrafish, ephrinb2 is required for preventing epithelial cells from rearranging once they have been restored in PE segmentation (Choe, 2015). We used an antibody of Ephb2 on control and *Foxi3*^−/−^ embryos to determine if its expression may be affected at E9.0. We did not observe a loss of Ephb2 expression between *Foxi3*^+/−^ and *Foxi3*^−/−^ embryos within the epithelial cells (Figure 6A and 6C). This indicates that Ephb2 does not function downstream of Foxi3, and may act in an independent pathway.

**Figure 6:**
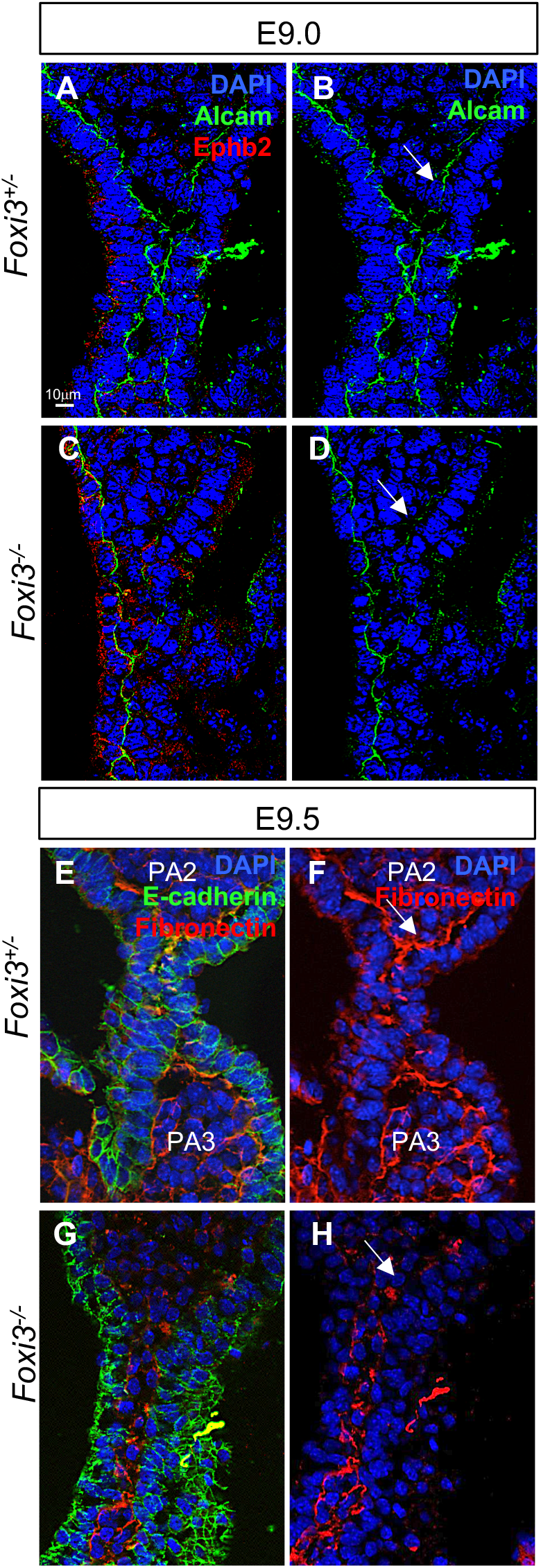
Alcam and Fibronectin expression is reduced in epithelial cells within the PA at E9.5 in *Foxi3*^−/−^ embryos. (A-D) DAPI (blue), Alcam (green) and Ephrinb2 (Ephb2, red) antibodies were used to examine embryo coronal sections at E9.0. Alcam localizes to the basal side of epithelium and Ephb2 localizes to the apical side. Sections from *Foxi3*^+/−^ control embryos are shown with all three fluorescence color channels in (A) and only DAPI and Alcam in (B). Sections from *Foxi3*^−/−^ embryos are shown with all three channels (C) and only DAPI and Alcam (D). White arrows indicate where Alcam expression is reduced in *Foxi3*^−/−^ mutants in comparison to control embryos (n=3). (E-H) DAPI (blue), E-cadherin (green) and Fibronectin (red) antibodies mark epithelial cells and the intracellular matrix that stabilizes the cells on coronal sections at E9.5. (E-F) Sections from *Foxi3*^+/−^ control embryos shown with all three fluorescent channels (E) and only the red and blue channels (F). (G-H) Sections from *Foxi3*^−/−^ embryos showing all three channels (G) and only the red and blue channels (H). White arrows indicate where expression is spotty and inconsistent in *Foxi3*^−/−^ mutants (n=3).

Fibronectin protein is an extracellular matrix protein that is present outside the basal surface of the epithelia closest to the adjacent mesenchyme and also expressed in the mesenchyme (Mittal et al., 2010; Trinh le, 2004). Fibronectin expression was absent or spotty adjacent to the endoderm and ectoderm in the mutant versus *Foxi3*^+/−^ control embryos (Figure 6E-6H). This indicates that *Foxi3* may have a role in regulating Fibronectin expression or localization. Together, this data confirms that epithelial cell morphogenesis affects adjacent Fibronectin expression.

## Discussion

In this report, we found that there is a genetic interaction between *Tbx1* and *Foxi3* in the formation of the thymus and parathyroid glands from PA3 in mammals. Inactivation of *Foxi3* in the *Tbx1* domain resulted in absent thymus and parathyroid glands. We then found that *Tbx1* and *Foxi3* are required autonomously in the PE for invagination to form pouches that marks the first transition in the epithelium in pouch formation. However, there are some differences in function of these two genes in the two dynamic transitions of the PE. We found that *Tbx1* is required autonomously in the first transition, to promote the formation of a stratified epithelium from an intercalated monolayer of cells, whereas *Foxi3* is required in the second transition, autonomously to convert the stratified epithelium back to a monolayer. Together the two genes add new genetic and mechanistic insights into transcription factors responsible for pharyngeal segmentation in mammals.

### Epithelial cells undergo dynamic transitions in the vertebrate PA

During vertebrate embryonic development, the segmentation of the PA is needed to create individual arches that later form derivative structures including the thymus and parathyroid glands (Graham et al., 2005; Shone and Graham, 2014). We revisited the process of pharyngeal segmentation to better understand the functions of *Tbx1* and *Foxi3*. Our data indicates that there are two major epithelial transitions required for morphogenesis, as shown in the model in Figure 7. In the first transition, epithelial monolayers become temporarily stratified to form multilayers during invagination. In the second transition, as invagination finishes, the cells revert back to monolayers forming a junction between the endodermal pouch and ectodermal cleft. One possibility during this process is that the cells lose their epithelial characteristics. Expression of E-cadherin is a marker for epithelial tight junctions. The cells maintained E-cadherin expression throughout the process, during both transitions (Figure 7). Another possibility is that some of the cells lose polarity. Expression of ZO-1 is a marker for apical/basal cell polarity. We found that apical/basal polarity is retained in the cells within the inner most cell layer facing the foregut and outer most cell layer of the embryo, but not in internal layers, as identified by ZO-1 expression (Figure 7). This indicates that the internal epithelial cells must temporarily lose specific aspects of tight junctions during this phase, perhaps to allow for stratification to take place.

**Figure 7:**
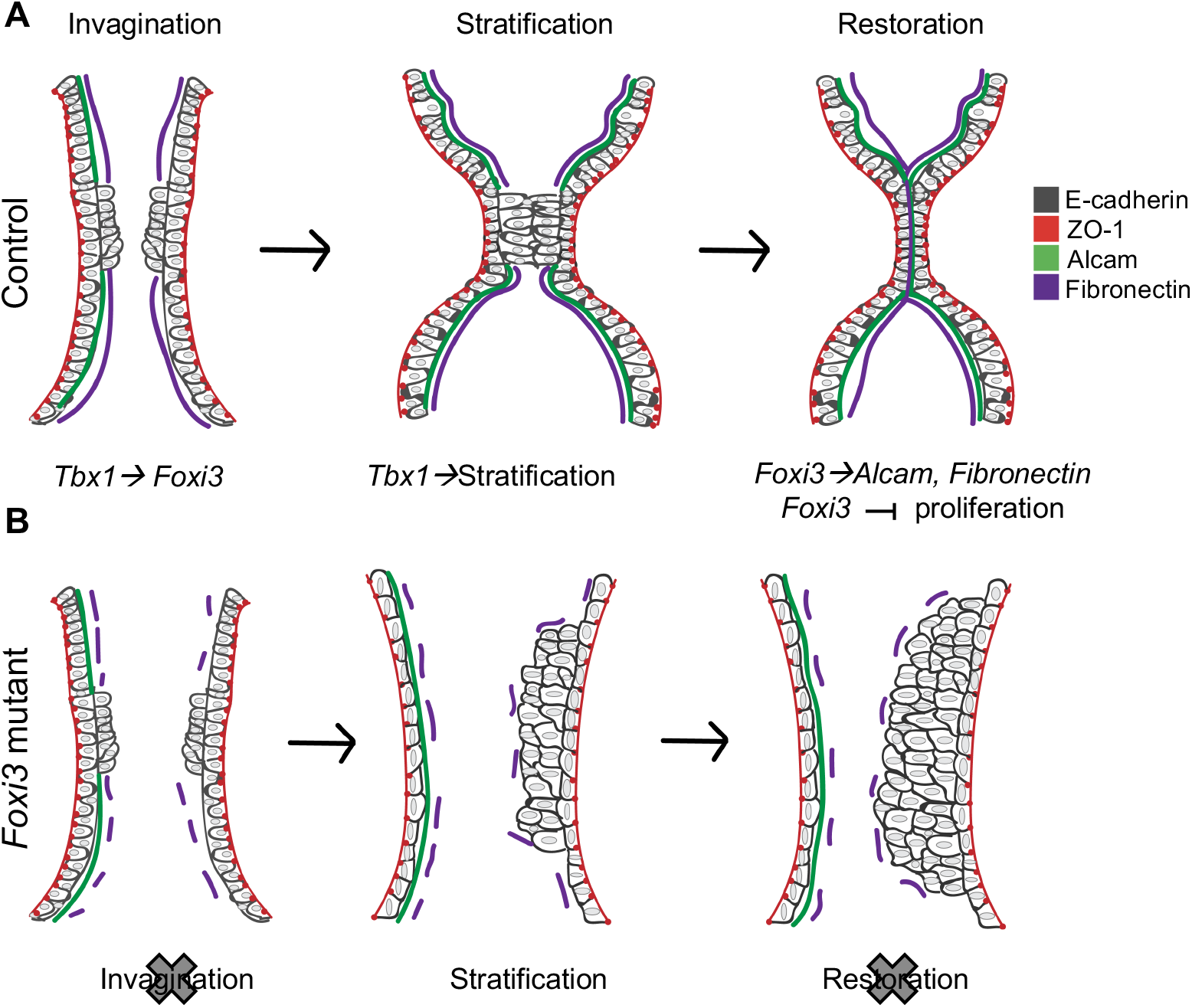
Summary cartoon of Tbx1 and Foxi3 functions in PA segmentation. (A) Cartoon of control epithelial cells during PA segmentation for each arch. E-cadherin is shown in green; ZO-1 is shown in red; Alcam is shown in blue and Fibronectin is shown in purple. When segmentation of an individual arch begins, the endoderm and ectoderm undergo invagination. *Tbx1* and *Foxi3* are required for invagination as indicated. At the same time as invagination, one or two additional layers of endoderm and ectoderm form in the position where the arch will segment. The outermost cell layer continues to maintain apical/basal polarity as depicted by ZO-1 expression. This is the first epithelial morphology transition. From examining of global *Tbx1* and PE mediated *Tbx1* conditional mutant embryos, *Tbx1* has an additional role as being required to form the transient multilayered epithelium. Alcam and Fibronectin are not expressed at the location of the stratified epithelium. After invagination is complete, an intercalated monolayer of cells are present at the junction of the endodermal pouch and ectodermal cleft. Alcam and Fibronectin are expressed throughout where segmentation has occurred. *Foxi3* acts upstream of Alcam and Fibronectin in the PE and inhibits cells proliferation during the stabilization process. (B) In *Foxi3*^−/−^ mutant embryos, invagination of endoderm and ectoderm is not initiated. The endoderm and ectoderm can form a transient stratified epithelium, but there is excess stratification of the PE to form many additional layers than the normal situation. Alcam is not expressed within the endoderm but is expressed within the ectoderm. Fibronectin expression spotty and is not continuous throughout the epithelia. The ectoderm in the *Foxi3* null mutant embryos have a more circular shape as opposed to a normal cuboidal shape.

A similar process has been described for branching morphogenesis to form pancreatic ducts from the distal foregut endoderm (Hick et al., 2009). In pancreatic organogenesis, a single layer of polarized foregut endoderm will invaginate into the mesenchyme to form branches and ducts composed of differentiated cells. In this process, a single layer of polarized epithelial cells is dynamically transformed to a multilayered epithelium, followed by a second transition back to a monolayer of polarized epithelial cells in the newly formed duct (Hick et al., 2009; Villasenor et al., 2010). As for the pharyngeal endoderm, the pancreatic endoderm cells express E-cadherin during this process and the row of cells on the apical side expresses ZO-1 (Hick et al., 2009; Villasenor et al., 2010).

A few years ago, a related process of dynamic transitions of the pharyngeal endoderm was described in zebrafish (Choe et al., 2013). During pharyngeal pouch formation, there is a similar two-step transition to a temporary stratified epithelium, which then becomes resolved, defined as stabilization of adherens junctions, in this experimental system, to two opposing layers when pouch formation becomes complete (Choe et al., 2013). There are several similarities between the zebrafish and mouse process, however, this has not yet been described in mammals. In zebrafish and mouse embryos, E-cadherin is expressed during both transitions. Further, polarity is lost within internal epithelial cells within the stratified endoderm (Choe et al., 2013; Choe and Crump, 2015). The focus on pharyngeal segmentation has been in defining the signaling mechanisms from the adjacent ectoderm and mesoderm to the PE. It was found that Wnt11r signaling emanating from the mesoderm is required to regulate destabilization of the endoderm to form transient multilayers (Choe et al., 2013). Independently, Wnt4a signals from the ectoderm upstream of Alcama (Alcam, mammals) expression and E-Cadherin, restore the PE to single cell layers with fortified tight junctions (Choe et al., 2013). Similarly, in mammals, we found that Alcam expression is disrupted in the region where the transient epithelial stratification takes place and becomes continuous after the second transition.

### Tbx1 functions to promote invagination and stratification cell autonomously

Previously, it was found in zebrafish and mice, that mesodermal *Tbx1* is required for pharyngeal segmentation. In zebrafish, the *tbx1* gene, in the mesoderm autonomously regulates expression of two important genes, *wnt11r* and *fgf8a* (Fibroblast growth factor 8a) that signals to the PE to promote pouch formation (Choe and Crump, 2014). Rescue of these defects took place when mesodermal *tbx1* expression was restored (Choe and Crump, 2014). Similarly, in mice, inactivation of *Tbx1* in the mesoderm results in a hypoplastic distal pharyngeal apparatus (Zhang et al., 2006) upstream of *Fgf8*. The function of *Tbx1* within the mesoderm, therefore is to signal to the endoderm to coordinate PA pouch formation (Choe and Crump, 2014). Thus, the data in zebrafish and mouse is consistent for a critical non-autonomous role of mesodermal *Tbx1* in pharyngeal segmentation.

One major difference between zebrafish and mice is in regards to *Tbx1* function in the PE. In zebrafish, the *tbx1* gene does not have an autonomous role in the PE in pouch formation (Choe and Crump, 2014). Further, in zebrafish, inactivation of *tbx1* did not affect expression of alcama in the PE cells (Choe and Crump, 2014). Global inactivation of *tbx1* and restoration in the PE failed to rescue pouch defects in zebrafish mutant larvae (Choe and Crump, 2014). In contrast, inactivation of *Tbx1* in the PE results in failed segmentation of the distal PA in mice (Arnold et al., 2006a; Jackson et al., 2014). Cell proliferation is not affected when *Tbx1* is inactivated in the endoderm using *Sox17^2A-iCre/+^*, but pouches do not form in these mutant embryos regardless (Jackson et al., 2014). In this report, we show that the PE cells fail to initiate the first transition of pharyngeal pouch formation, in both *Tbx1*^−/−^ and in the *Sox17^2A-Cre/+^;Tbx1^f/f^* mutant embryos. The cells retained expression of E-cadherin and ZO-1 indicating that, as expected, they did not lose their epithelial characteristics. This data indicates that *Tbx1* has a cell-autonomous role within the endoderm early during PA segmentation as shown in the model in Figure 7.

### Distinct roles of *Foxi3* in two phases of pouch epithelial transitions

In mammals, there are three *Foxi* class genes, *Foxi1, Foxi2* and *Foxi3*. In zebrafish, there is only one gene, termed *foxi1* and it has a similar expression pattern to that of *Foxi3* (Solomon et al., 2003). Recently, it was found that inactivation of *foxi1* resulted in a failure of the second transition in pharyngeal pouch formation in zebrafish (Jin et al., 2018). This is similar to our findings for *Foxi3* function in mammals, however we found that *Foxi3* is also important for the initial invagination of the epithelia. Further, in zebrafish, *foxi1* appears to have its major role in the ectoderm to signal non-autonomously to the PE and perhaps not autonomously in the PE (Jin et al., 2018). There is also a difference between zebrafish and mice versus mice regarding Foxi and Alcam expression. In the *foxi1* null mutant zebrafish, alcama expression appeared to be normal. In the mouse, loss of *Foxi3* resulted in failed or reduced Alcam expression, as shown in the model in Figure 7. It is possible that Alcam has a slightly different role downstream of *Foxi3* in mammals for this process.

Eph-ephrin signaling in adjacent cells, is important for cell migration. In zebrafish, EphB2 and EphB3 are required to maintain E-cadherin expression during budding morphogenesis of the endoderm from the foregut (Villasenor et al., 2010). During pharyngeal pouch morphogenesis in zebrafish, EphrinB signaling is required to increase expression of E-cadherin expression in the second transition (Choe, 2015). In *foxi1* mutant zebrafish, expression of EphrinB2a was not changed (Jin et al., 2018). Ephrin2a is expressed in a similar pattern to ZO-1. As in zebrafish, we found that EphrinB2a was not reduced in *Foxi3* mutant mouse embryos, and therefore is not directly implicated downstream of *Foxi3*.

In zebrafish, *foxi1* in the pharyngeal ectoderm initiates *wnt4a* signaling and that this is required for the second transition of the endoderm to form a final stabilized pouch (Jin et al., 2018). In PE specific knockout of *Foxi3* in mouse embryos, the ectoderm is able to invaginate and appears normal. However, if the main function of *Foxi3* is in the ectoderm, then we might have observed normal pouch development in the endodermal conditional mutant embryos. Our data also indicates that invagination of the ectoderm is not dependent on invagination of the endoderm. Rather, the ectoderm can initiate invagination independently. This is consistent with studies done in shark and chick embryos where the endoderm remains separate from the ectoderm throughout epithelial cell invagination (Shone and Graham, 2014).

### Role of Fibronectin and pharyngeal segmentation

Fibronectin is an extracellular matrix protein important as a substrate for cell migration that binds to the Integrin Beta 1 receptor in the epithelia (Mittal et al., 2010). These two proteins have important roles in epithelial and mesenchymal interactions needed for morphogenesis (Crump et al., 2004b; Mittal et al., 2010; Trinh le, 2004). Fibronectin and Integrins are important in development of neural crest cell (NCC) derived structures, such as the aortic arch arteries and aortic arch (Mittal et al., 2010). Here we found an essential role of *Foxi3* in the formation of arch arteries and the aortic arch, likely due to failed survival, migration and differentiation of NCCs (reviewed by (Mitchell et al., 2007). A previous study that focused on the function of *Foxi3* and craniofacial development, concluded that *Foxi3* promotes craniofacial NCC survival through FGF signaling from surrounding cells (Edlund et al., 2014). Our data suggests that disruption of Fibronectin expression could lead to failed NCC morphogenesis downstream of *Foxi3*.

Besides indirect functions on NCCs, it was found that Integrin Beta 1 in the endoderm forming the lung buds is required to prevent multilayering of the normal simple epithelium, and that when deleted, results in abnormal multilayer formation (Chen and Krasnow, 2012). Of interest, we found in normal morphogenesis that Fibronectin expression was disrupted during the first transition to form a transiently stratified epithelium, consistent with this possible normal function, and, as anticipated, it was restored during the second phase of PE transitions (Figure 7). In contrast, expression of Fibronectin was lower or spotty adjacent to the epithelia in global *Foxi3* null mutant embryos. It is possible that the endodermal cells are affecting Fibronectin expression, which then in turn allows more multilayers to form. Extensive multilayers formed only in the endoderm, but not the ectoderm, indicating that ectodermal cleft formation has separate requirements than that of the PE (Figure 7). This is similar to the situation with Alcam expression, where it was absent from the PE but still was present in the ectoderm in global *Foxi3* null mutant embryos (Figure 7). This underscores differences between the endoderm and ectoderm in PA segmentation.

### Translational Insights

Patients with 22q11.2DS have defects within structures derived from the PA including craniofacial, thymus aplasia, hypocalcemia, and aortic arch defects (McDonald-McGinn et al., 2015). *Tbx1* is among the deleted region within patients with this syndrome, and has been confirmed to have a role within the endoderm and mesoderm that are important for PA segmentation (Arnold et al., 2006b; Choe and Crump, 2014; Jackson et al., 2014). Based on our results, *Tbx1* may directly or indirectly regulate *Foxi3* expression that is also vital for normal development of the PA derived structures. There has been one report of a patient with a deletion of one allele of *Foxi3* that had severe ear defects, mild craniofacial defects, and arteries derived from PA1 and PA2 were absent (Tassano et al., 2015). These symptoms are different from the 22q11.2DS patients since the patient had normal heart, thymus, and parathyroid glands that are typically present in 22q11.2DS, however, it would be useful to consider variants in FOXI3 as a potential modifier of phenotype in 22q11.2DS.

## Materials and Methods

### Mouse mutant alleles

The following mouse mutant alleles used in this study have been previously described: *Foxi3^f/f^* (flox = f), *Foxi3*^+/−^(Edlund et al., 2014), *Tbx1^Cre/+^* (Vitelli et al., 2006) *Sox17^2A-iCre/+^* (Engert et al., 2009), *Tbx1*^+/−^ (Merscher et al., 2001) and *ROSA26^GFPf/+^* (RCE: loxP) (Sousa et al., 2009). *Foxi3*^−/−^ embryos were generated by inter-crossing *Foxi3*^+/−^ *Foxi3*^+/−^ mice. Double *Tbx1* and *Foxi3* heterozygous embryos were generated by inter-crossing *Tbx1*^+/−^ and *Foxi3*^+/−^ mice. The *Tbx1^Cre/+^;Foxi3^f/f^* and *Sox17^2A-iCre/+^;Foxi3^f/f^* embryos were generated by crossing male *Tbx1^Cre/+^;Foxi3^f/+^* or *Sox17^2A-iCre/+^;Foxi3^f/+^* mice with *Foxi3^f/f^* females. *Foxi3*^+/−^;*Tbx1^Cre/+^Foxi3^f/+^, Sox17^2A-iCre/+^;Foxi3^f/+^* and wildtype littermates were used as controls for the experiments.

The *Foxi3*^+/−^,*Sox17^2A-iCre/+^*, and *Tbx1^Cre/+^* mice were backcrossed 10 generations to a Swiss Webster background from a mixed C57Bl/6, Swiss Webster background. The PCR strategies for mouse genotyping have been described in the original reports and are available upon request. All experiments including mice were carried out according to regulatory standards defined by the NIH and the Institute for Animal Studies, Albert Einstein College of Medicine (https://www.einstein.yu.edu/administration/animal-studies/), IACUC protocol # 2016-0507.

### Mouse embryo heart histology and phenotypic analysis

Mouse embryos were isolated in phosphate-buffered saline (PBS) and fixed in 10% neutral buffered formalin (Sigma Corp.) overnight. Following fixation, the embryos were dehydrated through a graded ethanol series, embedded in paraffin and sectioned at 10 μm. All histological sections were stained with hematoxylin and eosin using standard protocols in the Einstein Histopathology Core Facility (http://www.einstein.yu.edu/histopathology/page.aspx). A total of 80 embryos, including controls, at E15.5 were obtained from more than 10 independent crosses and analyzed morphologically using light microscopy. Fisher’s exact test was used to determine if parathyroid and thymus defects were significant in *Tbx1*^+/−^; *Foxi3*^+/−^ compared to *Tbx1*^+/−^ embryos.

### RNAscope *in situ* hybridization

RNAscope *in situ* hybridization with non-radioactive mRNA probes was performed as previously described (Wang et al., 2012). Tissue was fixed in 4% paraformaldehyde (PFA) for 24 hours at 4°C and then cryopreserved in 30% sucrose in PBS overnight at 4°C. Embryos were embedded in OCT and cryosectioned at 10 μm. Probes for *Tbx1, Foxi3, Foxn1*, and *Gcm2* were generated by Advanced Cell Diagnostics.

### Whole mount *in situ* hybridization

Whole-mount RNA *in situ* hybridization with non-radioactive probes was performed as previously described (Alappat et al., 2005; Hidai et al., 1998), using PCR-based probes for *Foxi3* (Ohyama and Groves, 2004) and *Tbx1* (Funke et al., 2001). Following the whole mount protocol, the embryos were fixed in 4% PFA and then dehydrated through a series of graded ethanol, embedded in paraffin, and sectioned at 10 μm. Minimum of 3 embryos from 3 independent litters were analyzed for each experiment.

### Immunofluorescence on embryo sections

Embryos were collected at various stages: E8.5 (7-10 somite pairs), E9.0 (15-19 somite pairs), E9.5 (20-22 somite pairs), and E10.5 (31-33 somite pairs). Fixation was carried out in 4% PFA in PBS at 4°C for two hours. After fixation, tissue was washed in PBS and then cryoprotected in 30% sucrose in PBS overnight at 4°C. Embryos were embedded in OCT and cryosectioned at 10 μm. After fixation, frozen sections were obtained as described and then permeabilized in 0.5% Triton X-100 for 5 min. Blocking was performed with 5% goat serum in PBS/0.1% Triton X-100 (PBT) for 1 hour. Primary antibody was diluted in blocking solution and incubated for 1 hour. Primary antibodies used included: E-cadherin (BD Transduction laboratories 610181, 1:200 mouse), ZO-1 (Invitrogen 61-7300, 1:200 rabbit), Fibronectin (ab2413, 1:100 rabbit), Alcam (R&D BAM6561, 1:50 mouse), Ephrin b2 (ab150411, 1:200 rabbit) and GFP (ab6673 1:500 goat). Proliferation of cells was assessed by immunofluorescence using antibody anti-phospho Histone H3 (Ser10), a mitosis marker (06-570 Millipore). Sections were washed in PBT and incubated with a secondary antibody for 1 hour. Secondary antibodies were Alexa Fluor 488 goat a-mouse IgG (Invitrogen A32723) and Alexa Fluor 568 donkey a-rabbit IgG (Invitrogen A11019) at 1:500. Slides were mounted in hard-set mounting medium with DAPI (Vector Labs H-1500). Images were then captured using a Zeiss Axio Observer microscope with an apotome.

### Cell number and proliferation quantification on tissue sections

To count epithelial cell number, we obtained 10 μm serial coronal sections of control, *Foxi3*^−/−^, and *Sox17^2A-iCre/+^;Foxi3^f/f^* embryos, which were collected and stained with an antibody for E-cadherin. To ensure that the cell quantification was accurate, we counted E-cadherin positive cells in the PA in every other section throughout each embryo. We did not count epithelial cells that were not part of the PA. When counting cells, we matched the embryos by stage using somite counts, and we matched the sections by position within the embryo. We also ensured that for each pair of control and mutant embryos, we counted the same number of sections (10-12 per embryo). We counted all proliferating, phosphoH3 positive cells in each section and calculated the ratio of proliferating cells within the pharyngeal epithelium. Then, we estimated the mean and standard error of the average cell counts for controls and mutant embryos and compared them using the t-test. Representations of the complete PA region from at least 6 embryos per genotype from at least 3 independent litters were used in each assay.

## Acknowledgements, sources of funding and disclosures

We thank Silvia Racedo and Christopher De Bono for guidance and comments. We thank Hiroko Nomaru, Xia Wang, and Sophie Astrof for assisting with India ink injections. We thank Andrew Groves for *Foxi3* heterozygous and floxed mice and probes. We want to thank the Histopathology Facility at Einstein. This work was supported by NIH grant, P01HD070454 (BM), Leducq foundation network grant (BM and EH). The authors declare no conflict of interest.

## Author Contributions

Conceptualization, E.H. and B.E.M.; Methodology, Formal analysis, Investigation, Visualization, Writing-original draft and Project Administration E.H.; Writing—Review and editing, Supervision, and Funding aqusition, B.E.M.

## Supplementary Figure Legends

**Supplemental Figure 1: *Foxi3*^−/−^ and *Sox17-2A-iCre/+;Foxi3^f/f^* conditional mutants have PA defects and defects in PA derived structures.**

(A-C) Whole mount images of control (A), *Foxi3*^−/−^ (B), and *Sox17^2A-iCre/+^;Foxi3^f/f^* (C) embryos at E9.5. Asterisks indicate absent PA in both mutants.

(D-L) Transverse histology sections stained with H&E of control (D-F), *Foxi3*^−/−^ (G-I) and *Sox17^2A-iCre/+^;Foxi3^f/f^* (J-L) embryos at E15.5. Abbreviations: right subclavian artery (R. sub), thymus (Thy), aorta (Ao), pulmonary trunk (PT), right atria (RA), left atria (LA), right ventricle (RV), left ventricle (LV), ventricular septum (VS). Arrows indicate absent thymus glands (G, J), interrupted aortic arch type B (H, K), and ventricular septum defect (I).

**Supplementary Figure 2: *Tbx1*^+/−^;Foxi3^+/−^ have hypoplastic and ectopic parathyroid and thymus glands and *Tbx1^Cre/+^;Foxi3^f/f^* mutants have absent thymus and parathyroid glands.**

(A-F) Transverse histology sections stained with H&E of WT (A-B), *Tbx1*^+/−^;*Foxi3*^+/−^, (C-D), and *Tbx1^Cre/+^;Foxi3^f/f^* (E-F) embryos at E15.5. Abbreviations: parathyroid (PT), thyroid (T), thymus (Thy). Asterisks indicate absent thymus glands.

(G-I) H&E coronal histology sections of WT (G), *Tbx1*^+/−^;*Foxi3*^+/−^ (H), and *Tbx1^Cre/+^;Foxi3^f/f^*(I) embryos at E10.5. Arrows indicate the third pouch and morphology defect. Third pouch is smaller in (H) and absent in (I).

(J-L) RNAscope *in situ* hybridization with mRNA probes for *Gcm2* (green) and *Foxn1* (red) probes on sagittal sections in WT (J), *Tbx1^+/−^;Foxi3^+/−^* (K) and *Tbx1^Cre/+^;Foxi3^f/f^* (L) embryos at E11.5. *Gcm2* marks parathyroid precursor cells and *Foxn1* marks thymus precursor cells.

**Supplementary Figure 3: *Foxi3* expression is within the *Tbx1^Cre/+^ Foxi3^f/f^* and *Sox17^2A-iCre/^+;Foxi3^f/f^* lineages at E9.5 in comparison to controls.**

(A-H) WMISH with antisense *Foxi3* mRNA probes on whole mount embryos and coronal sections at E9.5. (A-B) *Sox17^2A-iCre/+^;Foxi3^f/+^* control, (C-D) *Sox17^2A-iCre/+^; Foxi3^f/f^* conditional mutant, (E-F) *Tbx1^Cre/+^;Foxi3^f/+^* control and (G-H) *Tbx1^Cre/+^; Foxi3^f/f^* conditional mutant whole mount embryos with corresponding coronal sections. Black arrows indicate where *Foxi3* mRNA expression is reduced within conditional mutants.

**Supplementary Figure 4: *Tbx1^Cre/+^;Foxi3^f/f^* embryos have no heart or arch artery defects.**

(A-F) Transverse histology sections stained with H&E of control (A-C) and *Tbx1^Cre/+^;Foxi3^f/f^* mutant (D-F) embryos. Abbreviations: right subclavian artery (R. sub), thymus (Thy), aorta (Ao), pulmonary trunk (PT), right atria (RA), left atria (LA), right ventricle (RV), left ventricle (LV), ventricular septum (VS). (G-H) India ink was injected into the ventricle of control (G) and mutant (H) embryos at E10.5 to visualize the aortic arches.

**Supplemental Figure 5: India ink injections reveals absent 4^th^ aortic arches in *Foxi3*^−/−^ and *Sox17^2A-iCre/+^;Foxi3^f/f^* mutant embryos at E10.5.**

(A-F) India ink was injected into the ventricle of control (A-B), *Foxi3*^−/−^ (C-D) and *Sox17^2A-iCre/+^;Foxi3^f/f^* (E-F) embryos at E10.5. The right and left side of these embryos are shown. Arrows indicate absent 4^th^ aortic arches in both mutants.

**Supplement Figure 6: Epithelial cell invagination is delayed in *Foxi3*^−/−^ and conditional mutant.**

(A-I) DAPI (blue), E-cadherin (green), and ZO-1 (red) antibodies were used to visualize epithelial cells within the PA at E10.5 on WT control (A-C), *Foxi3*^−/−^ (D-F), and *Sox17^2A-iCre/+^;Foxi3^f/f^*(G-I) coronal sections (n=4 for each control and mutant). White boxes indicate the location of higher magnification images.

**Supplemental Figure 7: Epithelial cell proliferation increases at E8.5 and not at E9.5 in *Foxi3* mutants.**

(A-L) Proliferation assay was performed using a phospho-H3 (pH3) antibody on coronal sections of WT control (A-D), *Foxi3*^−/−^ (E-F), and *Sox17^2A-iCre/+^;Foxi3^f/f^* (I-L) mutant embryos at E8.5-E9.5.

(M-N) Quantification of proliferation assays include counting E-cadherin positive epithelial cells within the PA and calculating the mitotic index. This is the ratio of pH3 positive cells within the E-cadherin positive epithelial cells. Error bars represent standard error, and asterisks indicate P value <0.05. Endoderm (End) and Ectoderm (Ect) were quantified separately in conditional mutants. At E8.5, a total of n=6 was done for each control and mutant embryo; at E9.5 a total of n=4 was done for each control and mutant.

